# Fluid-Structure Interaction Dynamics of Cervical Lymphatic Vessel Pumping and Valvular Function

**DOI:** 10.1101/2025.11.21.689846

**Authors:** Daehyun Kim, Jeffrey Tithof

## Abstract

A substantial portion of cerebrospinal fluid (CSF) drains through cervical lymphatic vessels (CLVs), a pathway mediated by basal and dorsal meningeal lymphatics. Impaired drainage along this route has been implicated in aging, Alzheimer’s disease, and traumatic brain injury. Despite considerable experimental investigation of CLV structure and function, computational modeling of this pathway remains limited. Here, we present a fully coupled two-dimensional fluid-structure interaction (FSI) model of a murine CLV constructed using the Lattice Boltzmann method for fluid dynamics and the immersed boundary method for the vessel geometry. Distinct from previous lymphatic vessel models, this framework is parameterized using data from recent in vivo imaging studies of CLVs. Using this model, we characterize the transient FSI dynamics within a single lymphangion, the pumping performance across a chain of three lymphangions with varying contraction phase delays, and the role of circular sinus geometry in regulating CSF transport under both favorable and adverse pressure gradients. Our results provide the first high-fidelity simulation of CSF drainage through CLVs, bridging a gap between experimental observations and mechanistic understanding. This work offers new insights into CLV pumping behavior and valve function, which helps inform the design of future experiments and therapeutic strategies aimed at enhancing CSF clearance.

## 1 Introduction

Historically, the brain was thought to be devoid of lymphatic vessels. This understanding was dramatically revised by the discovery of meningeal lymphatic vessels [1, 2], which facilitate cerebrospinal fluid (CSF) clearance. This pathway allows CSF to exit the brain, drain into the cervical lymphatic vessels (CLVs) [3, 4], and finally reach the cervical lymph nodes [5–7]. This CSF drainage route has gained significant attention due to its linkage with various neurological diseases. Diminished clearance of CSF may contribute to impaired brain function in the context of aging [8, 9], Alzheimer’s disease [9, 10], and other neurological conditions such as stroke [11] and traumatic brain injury [12, 13].

Largely due to the clinical associations, this CSF drainage route is a subject of active experimental investigation [14]. Many studies have focused on manipulating the meningeal lymphatic network to explore novel therapeutic possibilities [13, 15]. However, the complete network is still being mapped, as highlighted by the recent discovery of the nasopharyngeal lymphatic plexus [16] that is connected to the nasal lymphatics. Experimental approaches have targeted various components of the cervical lymphatic system. For instance, Jin et al. [17] demonstrated that CSF drainage could be enhanced by manipulating superficial lymphatic vessels using manual lymphatic drainage techniques. Du et al. [8] revealed that, in mice, deep cervical lymphatic vessels lose their pumping functionality with age. They further demonstrated that rescuing this pumping frequency by increasing smooth muscle contractility led to enhanced CSF clearance, suggesting a promising therapeutic target.

While there are numerous experimental studies on CSF drainage through the CLVs, there is a distinct lack of numerical simulation of this pathway. To date, few computational models have addressed this topic. Vinje et al. [18] formulated a simple yet comprehensive compartment model of CSF clearance pathways, which included various resistances representing routes such as the cribriform plate and perivascular spaces, to study CSF flow distribution under elevated intracranial pressure. Separately, our group previously developed a lumped parameter model representing the nasal lymphatics and CLVs that drain CSF [19]. To our knowledge, these are the only computational studies focused specifically on CSF drainage through the lymphatic system.

Despite limited modeling of CSF drainage, numerous prior numerical studies have investigated function of other peripheral lymphatic vessels. Indeed, the unique physiological features of lymphatic vessels, such as their intrinsic contractility and passive intraluminal valves (secondary valves), have attracted interest from engineers seeking to understand their transport mechanisms [20]. Bertram and Moore led much of this work with their foundational lumped parameter model (0D) of a lymphatic vessel segment [21–24], which also served as the basis for our group’s previous CLV model. The functional unit of a collecting lymphatic vessel is the lymphangion, defined as the segment of vessel between two adjacent intraluminal valves. This 0D framework has been widely used to investigate the intrinsic pumping behavior of lymphangions [21, 23], specifically how valve resistance [22], contraction frequency [25], transmural pressure [24], and lymphangion length govern lymph flow [22]. They have also been extended to explore the effects of phase delays between contractions and initial lymphatics branching effect on overall transport efficiency [26].

Beyond these 0D models, two-dimensional (2D) and three-dimensional (3D) simulations of lymphatic vessels have also been developed. Li et al. [27] developed a 2D Lattice Boltzmann model of a single lymphangion bounded by two valves, which explicitly coupled *Ca*^2+^-modulated active wall contractions with nitric oxide production driven by shear stress at the vessel walls and valve leaflets. They later expanded this model to a series of lymphangions [28], still including the ion coupling, to test the effect of gravity on lymphatic transport. Elich et al. [29] developed a 2D chain of contracting lymphangions using the immersed boundary method, which they used to explore lymphangion pumping direction (i.e., phase-delays between contractions) under various axial pressure differences, including adverse pressure drops. Wilson et al. [30] developed a 3D valve model to test how the circular sinus region enhances lymph transport under a positive pressure drop, ultimately seeking to determine the hydraulic resistance of the valve opening. This initial model was not a fully-coupled fluid-structure interaction (FSI) simulation; instead it used a sequential approach where computational fluid dynamics was used to estimate pressure applied to the valve. These pressures were then applied in a finite element analysis to estimate the valve deformation, followed by a final computational fluid dynamics run at that deformed position. Subsequently, they expanded this model to utilize fully-coupled FSI [31], enabling a more detailed investigation of the fluid dynamics and the resulting stress and deformation within the valve. In parallel, Ballard et al. [32] developed a 3D lymphatic valve model using the Lattice Boltzmann method and Lattice Spring methods. They found that valve stiffening could seriously impact lymphatic transport, a finding that supports recent experimental work [8]. They further expanded their model to incorporate both peristaltic vessel wall pumping and intervening valves to determine optimal pumping dynamics [33]. Adeli Koudehi et al. [34] developed a 3D FSI model of murine collecting lymphatic vessels where valves were designed as porous membranes whose state (open or closed) was determined by their permeability, an approach similar to that of Bertram’s work [21].

It is worthy to note that the majority of the lymphatic simulations previously described are based on parameters derived from rat mesenteric lymphatics [35–37]. Consequently, advanced computational models focusing purely on the unique dynamics of CLVs are notably lacking. While the overall mechanisms governing lymph flow may be similar between these regions and species, key geometric properties (e.g., diameter, lymphangion length) and fluid dynamic parameters (e.g., Reynolds and Womersley numbers) are expected to differ. Furthermore, given the rising research interest in CLVs in the context of neurological disease, there is a clear need for computational studies that can investigate parametric sensitivity that is difficult or impossible to quantify experimentally (e.g., the effects of varied inter-lymphangion contraction phasing, manipulations of parameters). While our group’s previous 0D model captured the CSF drainage pathway through the CLVs, it was not designed to resolve the transient flow dynamics within the CLVs themselves.

In this study, we developed a new 2D CLV model that leverages fully-coupled FSI. We use the Lattice Boltzmann method to solve the fluid dynamics, coupled with the immersed boundary method to model the CLV, including the vessel wall motion and passive valves. Critically, the model adopts parameters from murine CLVs based on imaging from recent experimental studies [8, 12, 17]. Using this model, we investigate the detailed FSI dynamics within a single lymphangion, evaluate the pumping efficiency by varying the contraction phase between three lymphangions, and assess the lymphatic transport efficiency for different circular sinus bulb geometry at the valves for both positive and negative pressure difference across the valves. This study provides a deeper understanding of CSF drainage through CLVs, and it offer new insights relevant for design of therapeutic devices that enhance CSF drainage.

## 2 Methods

### 2.1 Lattice Boltzmann Method

We use the Lattice Boltzmann method (LBM) to simulate the transient fluid flow, owing to its computational efficiency. The LBM is based on the equation:

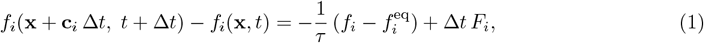

where *f*_*i*_ represents the particle distribution along discrete velocity **c**_*i*_, Δ*t* is the time step (typically Δ*t* = 1 in lattice units), *τ* is the relaxation time, *F*_*i*_ is the discrete forcing term, and 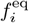 is the local Maxwell-Boltzmann equilibrium. For an isothermal lattice with a speed of sound 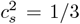, this equilibrium is computed as:

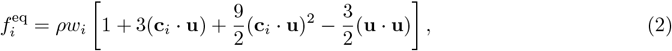

where *ρ* is the fluid density and **u** is the macroscopic fluid velocity.

The discrete force *F*_*i*_ incorporates the effect of the macroscopic force density **F** via Guo’s forcing scheme [38]. Specifically, the collision step is expressed as:

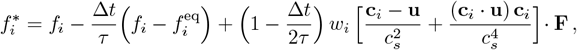

where *w*_*i*_ is the weight coefficient for the D2Q9 lattice. The streaming step is then 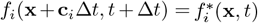 . After streaming, the macroscopic density and momentum are obtained by:

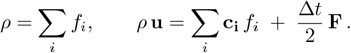

A more detailed D2Q9 mathematical formulation and its implementations can be found in the textbook [39], which also served as a basis of this work.

### 2.2 Immersed Boundary Method

The immersed boundary (IB) method [40] is used to model the CLV geometry immersed in the fluid, coupling the fluid on an Eulerian grid (Ω) with the CLV on a Lagrangian grid (Γ). The macroscopic force density (**F**(**x**, *t*)), which determines the discrete forcing term *F*_*i*_ in Eq. (1), is computed by spreading the Lagrangian force:

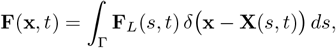

where **F**_*L*_ is the Lagrangian force density, *s* is the curvilinear material coordinate, and **X**(*s, t*) is physical spatial coordinate of the material point *s* at time *t*. The function *δ* is a Dirac delta function that acts as the kernel for interpolation between the Eulerian and Lagrangian grids. We employ the four-point kernel *ϕ*(*r*) based on Peskin’s original work [40]:

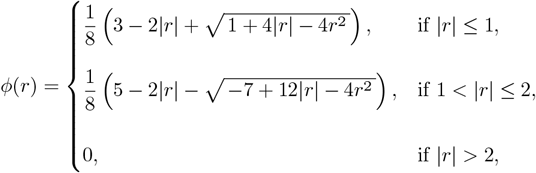

where *r* represents the normalized distance between Eulerian grid point (fluid node) and Lagrangian marker. The two-dimensional discrete delta function *δ* is then obtained as a separable product of the one-dimensional kernels:

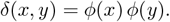

To update the position of the Lagrangian points due to forcing from the fluid, the Eulerian fluid velocity is interpolated to the Lagrangian grid using the same delta function kernel:

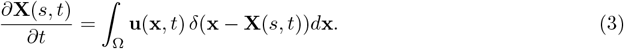

A key aspect of the IB method lies in formulating **F**_*L*_. We adopt a spring-based model to construct these Lagrangian forces. In this work, the CLV is modeled as a combination of static walls, moving walls, and valve components.

#### 2.2.1 Static walls

The static walls are modeled to represent the circular sinus regions near the valve and the walls at the inlet and outlet boundaries. These components enforce a zero-velocity condition at the Lagrangian nodes by applying a velocity-penalty formulation:

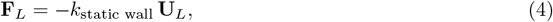

where *k*_static wall_ is the penalty stiffness coefficient and **U**_*L*_ is the local Lagrangian velocity interpolated from the fluid field. This penalty force resists the motion of the IB, effectively enforcing the near no-slip condition with negligible displacement of the wall.

#### 2.2.2 Pumping wall

For the lymphangion walls, a prescribed motion is defined to enforce intrinsic pumping. A spring-damper formulation [41] is used to ensure the Lagrangian points follow this motion while enforcing the near no-slip condition:

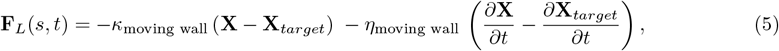

where *κ*_*moving wall*_ is the stiffness coefficient and *η*_*moving wall*_ is the damping coefficient. The term **X**_*target*_ denotes the prescribed target position for the curvilinear material coordinate *s*, which represents a one-dimensional Lagrangian coordinate along the vessel wall (see Figure 1), at time *t*. The target geometry **X**_*target*_ is defined by superimposing a time-varying vertical displacement (*y*_*target*_ ∈ **X**_*target*_) onto a fixed reference shape, represented by *y*_*linear*_:

**Fig. 1.**
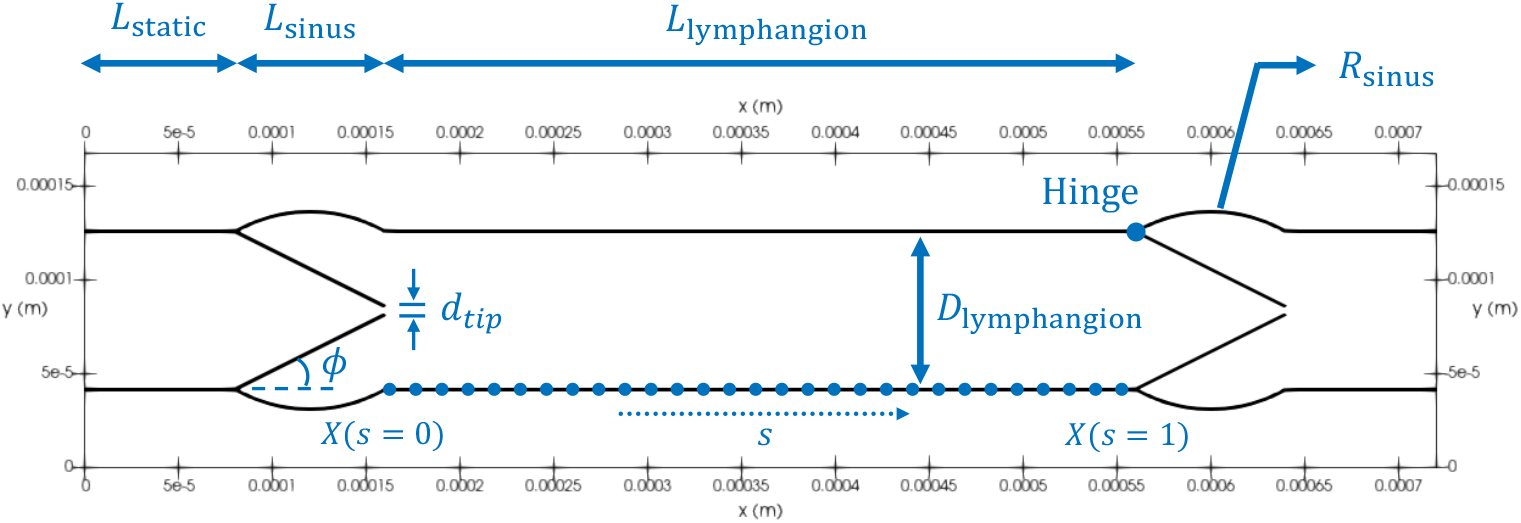
Schematic of the 2D computational domain for a single lymphangion. The model consists of a static inlet region (*L*_*static*_), an inlet valve as well as its sinus (*L*_*sinus*_), a central pumping chamber (*L*_*lymphangion*_) bounded by valves. Key geometric parameters, including vessel diameter (*D*_*lymphangion*_), valve gap (*d*_*tip*_), and sinus curvature (*R*_*sinus*_) are labeled, and the valve hinge location is explicitly marked. The curvilinear Lagrangian coordinate *s* is illustrated along the lower wall, bounded by the non-dimensional material coordinates *s* = 0 and *s* = 1. Lagrangian markers (dots) are illustrated only in this segment to improve visual clarity, while the entire vessel geometry is modeled using the immersed boundary method.

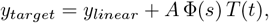

where *A* is the contraction amplitude, Φ(*s*) is a spatial envelope function, and *T* (*t*) is a temporal waveform defining the contraction cycle.

More specifically, Φ(*s*) is a piecewise function that defines the spatial profile of the contraction:

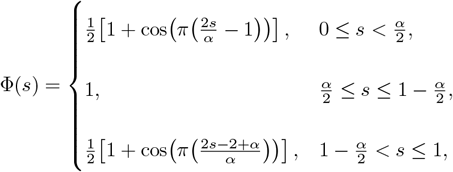

and *T* (*t*) is the asymmetric temporal waveform that drives the motion:

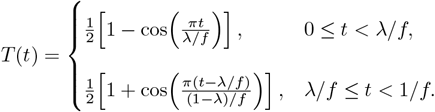

Here, *f* is the contraction frequency and *λ* is the duty ratio. The Φ(*s*) function ensures that both ends of the lymphangion wall remain fixed (*s* = 0, *s* = 1, non-dimensional) while the most central region undergoes a spatially uniform contraction. The parameter *α* controls the width of the tapering regions, thereby defining the width of the uniform central region as 1−*α*. The asymmetric waveform *T* (*t*) produces a faster contraction phase (duration *λ/f*) and a slower relaxation phase (duration (1 − *λ*)*/f*) [20].

Figure 2 helps to visualize these functions. Figure 2A illustrates representative profiles of Φ(*s*) for different values of *α*. As *α* increases, the width of the central, uniformly contracting region (1 − *α*) becomes narrower. Figure 2B illustrates the asymmetric contraction in time, where a value of *λ* = 0.5 results in a symmetric cycle.

**Fig. 2.**
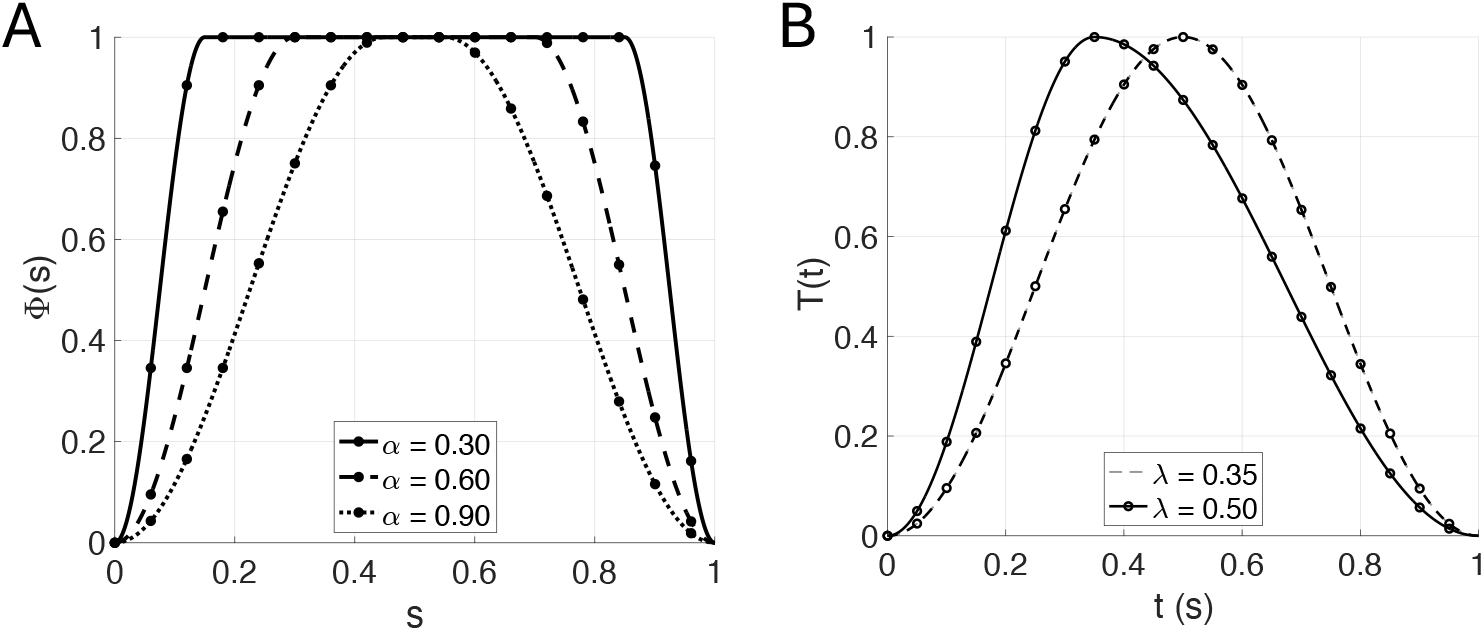
Illustrations of the spatial and temporal functions used to define the pumping wall motion in a single lymphangion. The spatial envelope function Φ(*s*) for three different tapering parameters: *α* = 0.30 (solid), *α* = 0.60 (dashed), and *α* = 0.90 (dotted). The parameter *α* controls the width of the tapering regions, thus defining the central uniformly contracting zone (1 − *α*). Note that the tip of Valve 1 is located at *s* = 0 initially, and the hinge of Valve 2 is located at *s* = 1. (B) The temporal waveform *T* (*t*) for a single cycle. The symmetric case (*λ* = 0.5) is shown for comparison against the asymmetric case (*λ* = 0.35), which models the characteristic fast contraction and slow relaxation behavior.

In our simulation, we set *α* = 0.3, which defines a central fully-contracted zone spanning 70% of the lymphangion length. We set *λ* = 0.35 to model the characteristic fast-contraction, slower-relaxation behavior observed experimentally [4].

#### 2.2.3 Valves

Each bileaflet valve (secondary valves in the lymphatic system) is modeled as a rigid body that rotates passively about a hinge under the influence of the surrounding fluid [42–44]. At every Lagrangian node, a linear penalty force is applied to enforce the rigid-body motion:

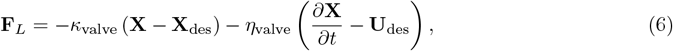

where *κ*_valve_ is the penalty coefficient and *η*_valve_ is the damping coefficient. The variable **U**_des_ = ***ω*** × **r** is the desired rigid-body velocity, ***ω*** is the angular velocity, and **r** is the position vector of the Lagrangian point relative to the hinge. The location of the hinge is illustrated in Figure 1.

The net hydrodynamic torque acting on the leaflet is obtained by integrating the moment of the Lagrangian forces:

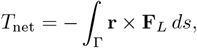

from which the angular acceleration is computed as 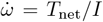, with *I* being the moment of inertia about the hinge. The angular velocity *ω* and the rotation angle *ϕ* are then updated explicitly in time, subject to angular limits (*ϕ*_*min*_ and *ϕ*_*max*_) that define the allowable range of motion. The values of *ϕ*_*min*_ and *ϕ*_*max*_ are provided in Table 1.

**Table 1.**
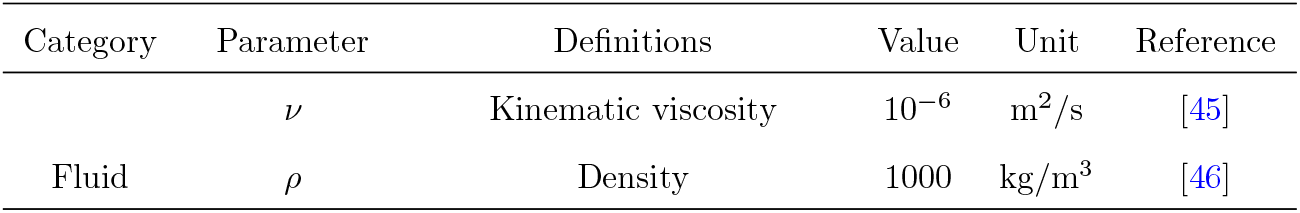
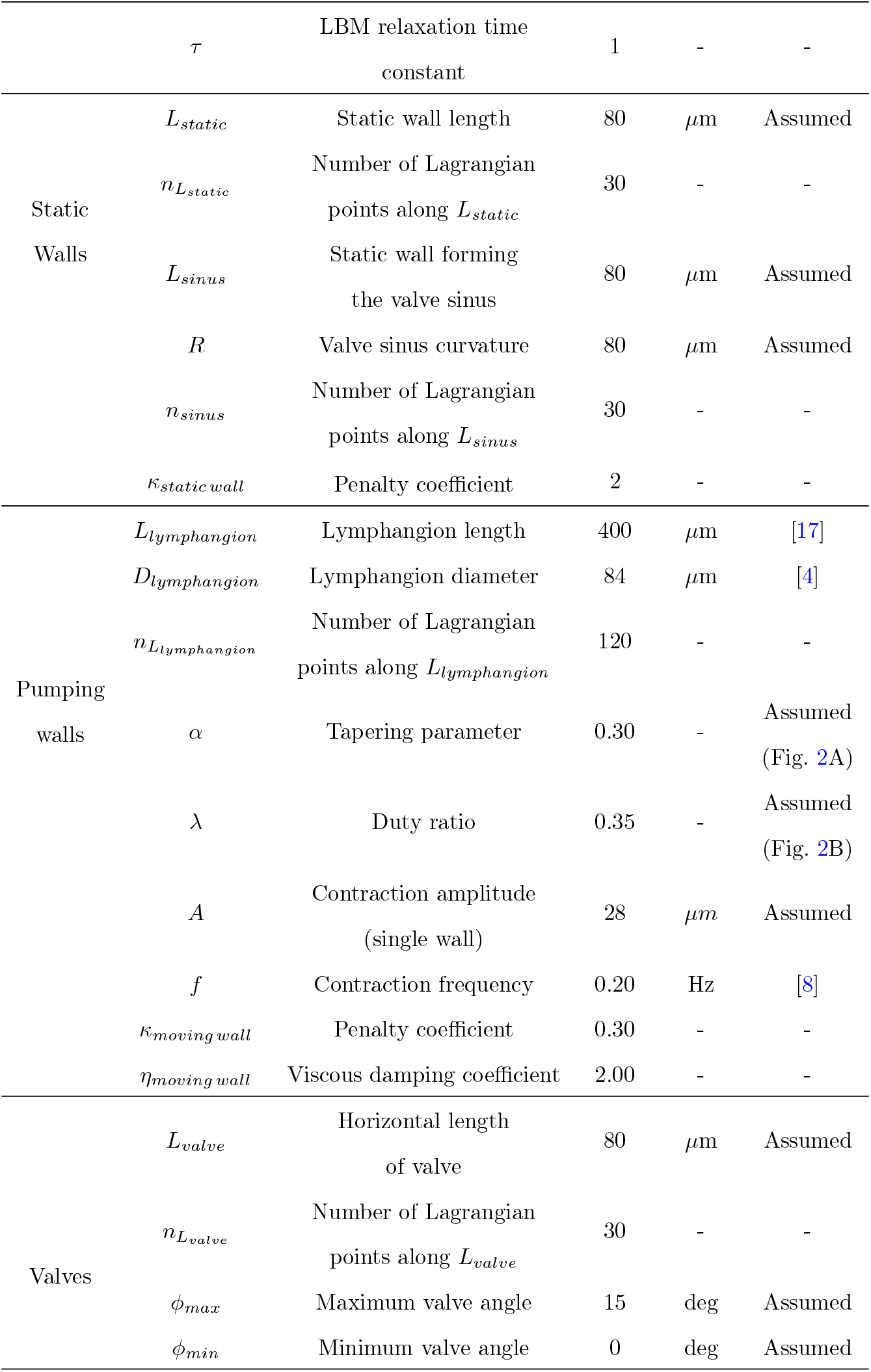
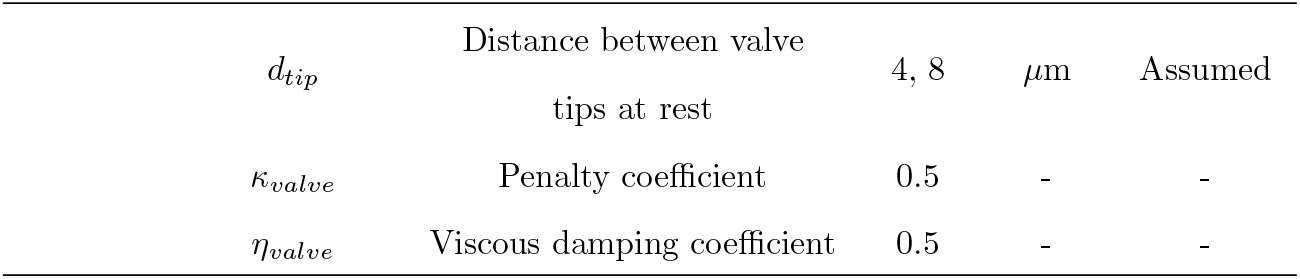
Summary of model parameters, including fluid properties, lymphangion geometry, and IB coefficients.

Finally, the desired leaflet geometry **X**_des_ is updated based on the new angle:

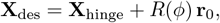

where *R*(*ϕ*) is the rotation matrix for angle *ϕ* and **r**_0_ is the initial position vector of each node relative to the hinge. This formulation provides two-way fluid-structure coupling: fluid-induced torques drive the leaflet rotation, while the resulting motion alters the surrounding flow through the IB forcing. This model assumes the leaflet is rigid and does not deform.

#### 2.2.4 Numerical setup

The geometry of a single lymphangion is illustrated in Figure 1. The model consists of a static inlet region (*L*_*static*_), an inlet valve within its sinus (*L*_*sinus*_), a central pumping chamber (*L*_*lymphangion*_), an outlet valve and sinus, and a static outlet region. The specific parameters for this geometry are listed in Table 1.

The simulation domain’s total height is set to twice the central lymphangion diameter (*D*_*lymphangion*_), with the vessel positioned at the vertical center. The fluid, representing a mixture of CSF and interstitial fluid, is modeled using the properties of water [45, 46]. The far inlet and outlet walls (with length *L*_*static*_) as well as the curved valve sinus (with curvature *R*_*sinus*_) are treated as static walls, inspired by CLV geometries observed in Jin et al [17]. The lymphangion is not modeled as a perfectly closed system; at rest, a non-zero gap, *d*_*tip*_, exists between the valve leaflet tips. The simulation was implemented in Python, primarily utilizing the Numpy [47] and Numba [48] libraries for performance. All the post-processing was performed in ParaView [49].

For the single and three lymphangion studies, a zero-pressure (*P* = 0 *Pa*) condition was applied at the far left and right boundaries of the domain to isolate the flow and ensure it is driven purely by the pumping and valve motions. A periodic boundary condition was applied to the top and bottom boundaries, which is a common practice in IB simulations to simulate an unbounded fluid environment. The LBM-IB solver was verified against the analytical solution for Hagen-Poiseuille flow (a steady, laminar flow in a channel driven by a prescribed pressure gradient). This verification is detailed in Appendix A. Our simulation shows a good agreement with the analytical solution, with a relative error in the maximum velocity of approximately 0.3 %. A grid convergence study was also performed for the single lymphangion case. We examined how the time-averaged outlet volumetric flow rate (⟨*Q*_*out*_⟩_*t*_) varied with the fluid grid refinement, while all IB parameters were held constant. This grid convergence study is detailed in Appendix B. The relative error in ⟨*Q*_*out*_⟩_*t*_ between the two finest grids tested (67, 000 and 94, 560 cells, respectively) was approximately 5.7 %. To balance solution accuracy and computational efficiency, the 67, 000-cell grid was selected for all subsequent simulations, as the computational cost increases exponentially with finer discretizations.

## 3 Results

### 3.1 Pumping Dynamics of a Single Lymphangion

We first discuss the key fluid-structure interaction dynamics of the single lymphangion model simulated over two pumping cycles, as shown in Figure 3. The prescribed motion of the pumping wall is shown in Figure 3B, where the diameter oscillates between 84 *µ*m (diastole) and a minimum of 38 *µ*m (systole).

**Fig. 3.**
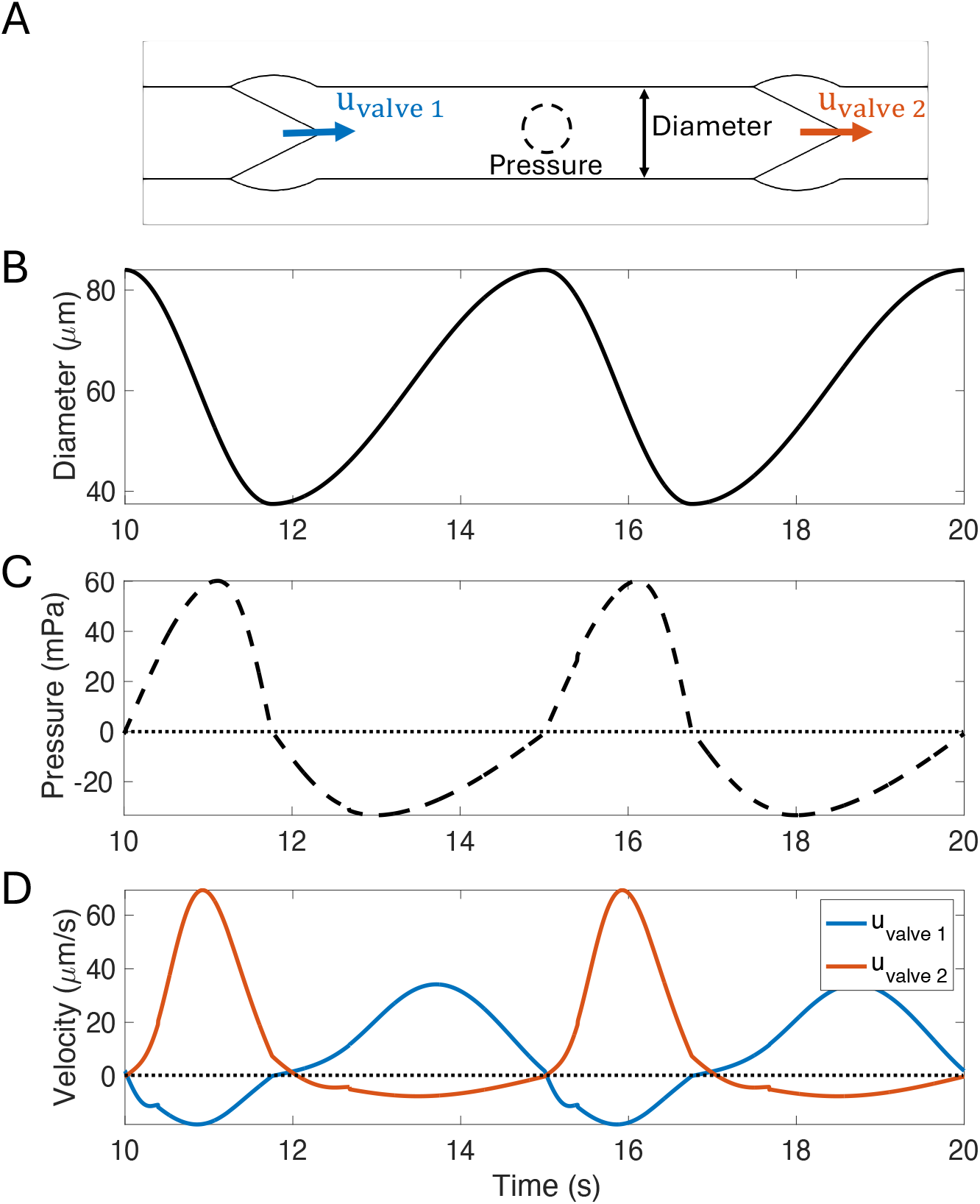
Pumping dynamics of the single lymphangion model over two cycles. (A) Schematic of the lymphangion indicating the locations for measuring wall diameter, fluid pressure (at the center), and axial velocities at the upstream valve (valve 1) and downstream valve (valve 2). (B) Prescribed wall diameter, varying from a maximum of 84 *µm* in diastole to a minimum of 38 *µm* in systole. (C) Resulting relative pressure in the center of the lymphangion. (D) Axial velocity at valve 1 and valve 2. Contraction induces significant forward flow at valve 2 and some backflow at valve 1.

The resulting relative pressure generated within the lymphangion chamber is shown in Figure 3C. A temporal offset between diameter and pressure is observed: the pressure reaches its maximum (∼ 50 mPa) mid-contraction at 11.8 s as the fluid is being forcefully expelled. Conversely, a significant negative pressure is generated as the chamber walls expand, creating suction to draw fluid in from the inlet.

This pressure differential drives the fluid flow, as shown by the axial velocities at the upstream valve (*u*_*valve* 1_) and downstream valve (*u*_*valve* 2_) in Figure 3D. During the contraction phase (e.g., 10–11.8 s), the rise in internal pressure causes significant forward flow (positive velocity) through the valve 2, while simultaneously forcing the valve 1 shut and causing a non-negligible backflow (negative velocity). This phase-dependent, valve-mediated dynamic is the core mechanism of CLV transport.

The small, sharp spikes visible in the velocity curves (e.g., at 10.5 s and 15.5 s in the valve 1 data and 12.5 s and 17.7 s in the valve 2 data) are minor numerical artifacts. They occur at the moment the rigid valve leaflet model reaches its hard-lock angular limit (*ϕ*_*min*_), causing a small, instantaneous disturbance in the flow field.

### 3.2 Pumping in a Three-Lymphangion Chain

To investigate the effect of coordinated pumping, we simulate three identical lymphangions in series. Figure 4A provides a schematic of the three-lymphangion model, defining the contraction phase delay Δ*Φ* between adjacent lymphangions and the measurement location for the outlet volumetric flow rate, *Q*, where

**Fig. 4.**
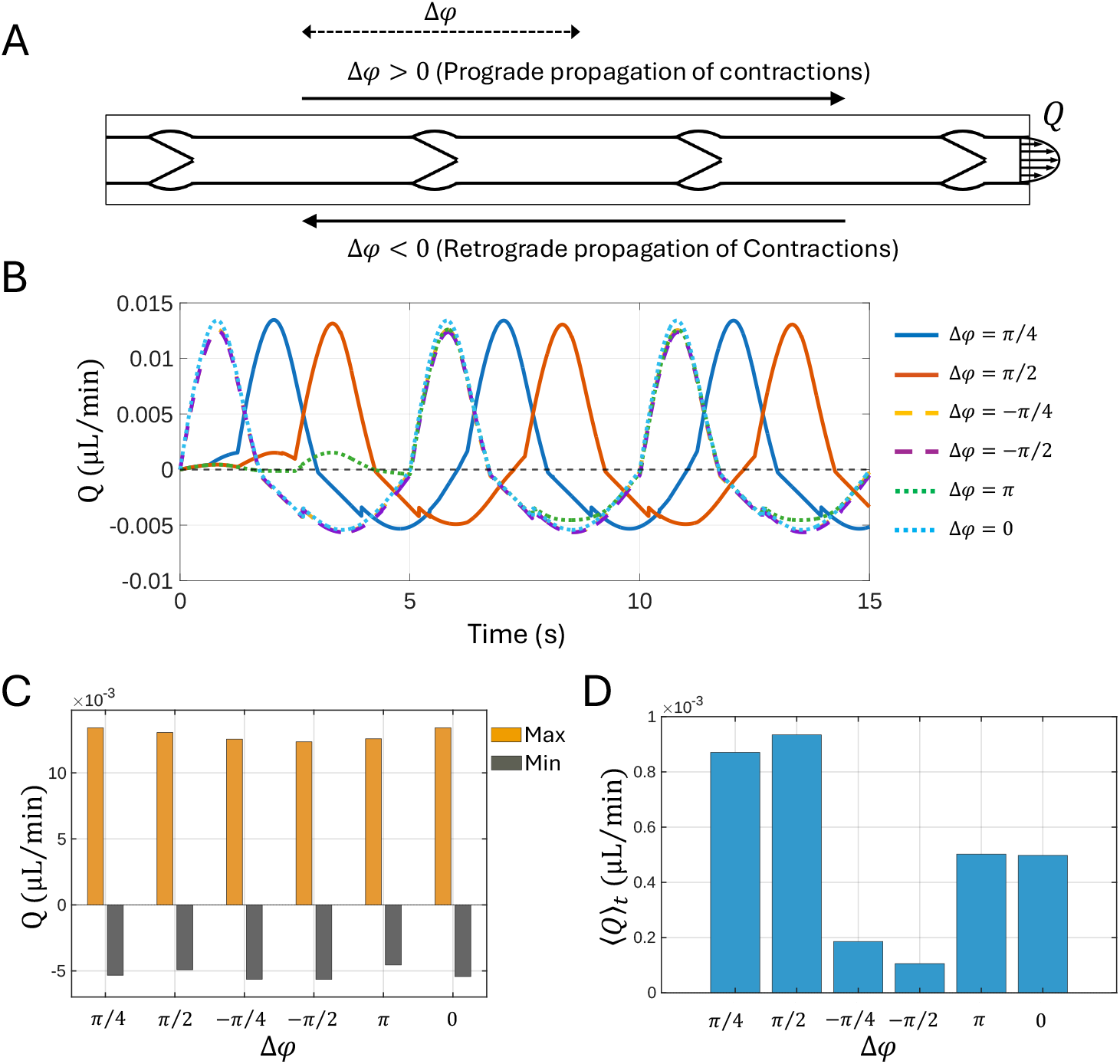
Pumping efficiency of a three-lymphangion chain under six different inter-lymphangion contraction phase delays (Δ*Φ*). (A) Schematic of the three-lymphangion model, defining the direction of prograde/retrograde contraction waves, the phase delay Δ*Φ*, and the measurement location of volume flow rate *Q*. (B) *Q* for all six phase-delay cases in time series. (C) Instantaneous maximum (forward flow) and minimum (backflow) volumetric flow rates extracted from (B). (D) Time-averaged mean outlet flow rate ⟨*Q*⟩_*t*_, indicating the net pumping efficiency for each case, extracted from (B).

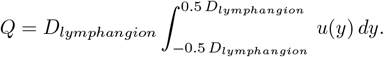

We tested how the *Q* is affected by the contraction phase delay between adjacent lymphangions. Six cases were simulated: prograde propagation of contractions (contraction propagating from left to right with Δ*Φ* = *π/*4 or Δ*Φ* = *π/*2), retrograde propagation of contractions (contractions propagating from right to left with Δ*Φ* = *π/*4 or Δ*Φ* = *π/*2), an alternating pattern (Δ*Φ* = *π*), and a synchronous contraction (Δ*Φ* = 0).

Figure 4B shows the time series of *Q* for all six cases, which exhibit a representative pattern: a strong forward flow pulse during contraction of the third lymphangion, followed by non-negligible backflow as the walls relax. The small, sharp bumps (e.g., at 4 s, 9 s, and 14 s in the propagating Δ*Φ* = *π/*4 case) are numerical artifacts from the valve leaflets reaching their hard-lock closed angles.

To compare performance, the instantaneous maximum, minimum, and time-averaged mean volume flow rates were computed (Figure 4C-D). These representative statistics were calculated only after the second contraction cycle of the third lymphangion (not shown in the figure), ensuring the system had reached a cyclical steady state. As shown in Figure 4C, the largest instantaneous forward flow (Max) was achieved by the prograde propagating wave with Δ*Φ* = *π/*4 and the synchronous case (Δ*Φ*=0). Conversely, the retrograde waves (e.g., Δ*Φ* = −*π/*4) generated the largest magnitude of instantaneous backflow (Min). The lowest magnitude of instantaneous backflow was produced by the alternating contraction (Δ*Φ* = *π*). For the prograde propagation waves, shorter phase delays (e.g., *π/*4 vs. *π/*2) produced higher instantaneous forward flow but also higher instantaneous backflow.

Figure 4D summarizes the net transport efficiency via the time-averaged flow, ⟨*Q*⟩_*t*_. The highest ⟨*Q*⟩_*t*_ was achieved by the propagating contraction wave with a phase delay of Δ*Φ* = *π/*2, which was slightly more effective than Δ*Φ* = *π/*4. The lowest ⟨*Q*⟩_*t*_ was produced by the retrograde propagation contraction waves, with Δ*Φ* = −*π/*2 resulting in the worst performance. The optimal propagating case (Δ*Φ* = *π/*2) produced a mean flow of 5.2 × 10^−4^ *µ*L/min, which was approximately 9.3 times larger than the 0.56 × 10^−4^ *µ*L/min produced by the worst case (Δ*Φ* = −*π/*2).

### 3.3 Influence of Sinus Geometry on Valve Dynamics

Recent high-resolution imaging of CLVs consistently reveals a rounded, bulbous sinus region in the vicinity of the valves [8, 16, 17]. Although the exact morphology of this sinus varies and is rarely perfectly circular, a rounded profile is characteristic. To isolate the effect of the valve sinus geometry, we simulated the valve’s opening dynamics in response to a constant 1 *Pa* positive pressure drop. We compared two geometries: a straight sinus (*R* = ∞) and the circular sinus (*R* = 80 *µm*). Both valves were initialized in a fully closed state. The results are shown in Figure 5. Figure 5A illustrates the flow driven by the positive pressure drop for both the straight and circular sinus geometries at a series of three snapshots. The velocity vectors are also shown, indicating a fully laminar flow profile as the valves open. The volumetric flow rate downstream, *Q*, is indicated in the figure, which was computed 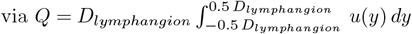.

**Fig. 5.**
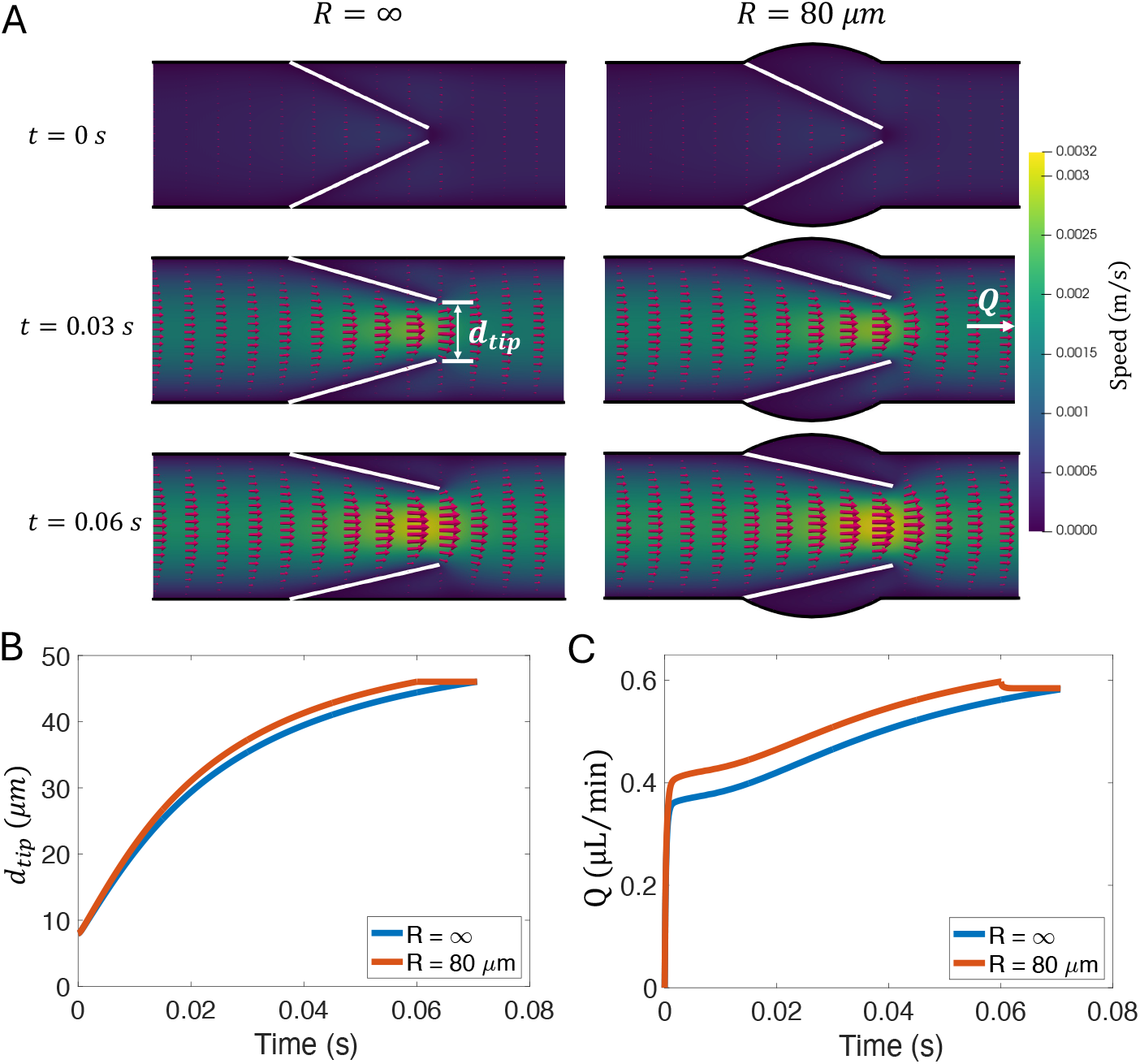
Valve opening dynamics under a constant 1 *Pa* positive pressure drop, comparing a straight sinus (*R* = ∞) with a circular sinus (*R* = 80 *µ*m). (A) Snapshots of the transient flow across the valve at *t* = 0, 0.03, and 0.06 s. (B) Time-series of the distance between valve tips (*d*_*tip*_). The valve with circular sinus case (orange curve) opens faster. (C) Comparison of the volumetric flow rate (*Q*) at the outlet.

The width of the valve opening is explicitly compared in Figure 5B, which plots the distance between the valve tips (*d*_*tip*_) over time. It was found that the valve with the circular sinus opened faster than the straight sinus, reaching its fully-opened state at approximately *t* = 0.06 *s*, while the valve in the straight sinus was still in the process of opening. The valve in the straight sinus required 0.07 s to fully open; this duration was set as the integration period, *T*_*open*_, for further analysis.

Figure 5C plots the volumetric flow rate at the downstream outlet. Since the valve with the circular sinus opens faster, it causes a higher overall instantaneous *Q*. The time-averaged volumetric flow rate 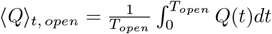 was computed for both cases over an integration period of *T*_open_ = 0.07 s. This calculation resulted in 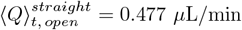 min versus 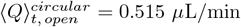. This 7.97% difference confirms that the circular sinus geometry enhances the valve opening and thus increases the volumetric flow rate.

A noticeable dip is visible in the circular sinus volumetric flow rate data (orange curve) at *t* ≈ 0.06 s. This corresponds to the moment the valve reaches its maximum angle (*ϕ*_*max*_) and abruptly stops opening as seen in Figure 5B. While the valve’s motion is purely passive, it does not open at a constant pace (Figure 5B); this evolving motion modestly contributes to driving the flow. Consequently, when this motion abruptly ceases, a small, transient decrease in the volumetric flow rate appears (Figure 5C). We next tested whether the circular sinus also helps close the valve more effectively. A negative pressure drop (-1 Pa) was prescribed across the axial domain while the valves were initially in a fully-opened state for both sinus cases. The results are shown in Figure 6. The velocity snapshots (Fig. 6A) illustrate the backflow driven by the negative pressure drop for both geometries. It is noteworthy that the overall closing time (less than 0.04 s) is shorter than the opening time observed in the previous test (0.07 s).

**Fig. 6.**
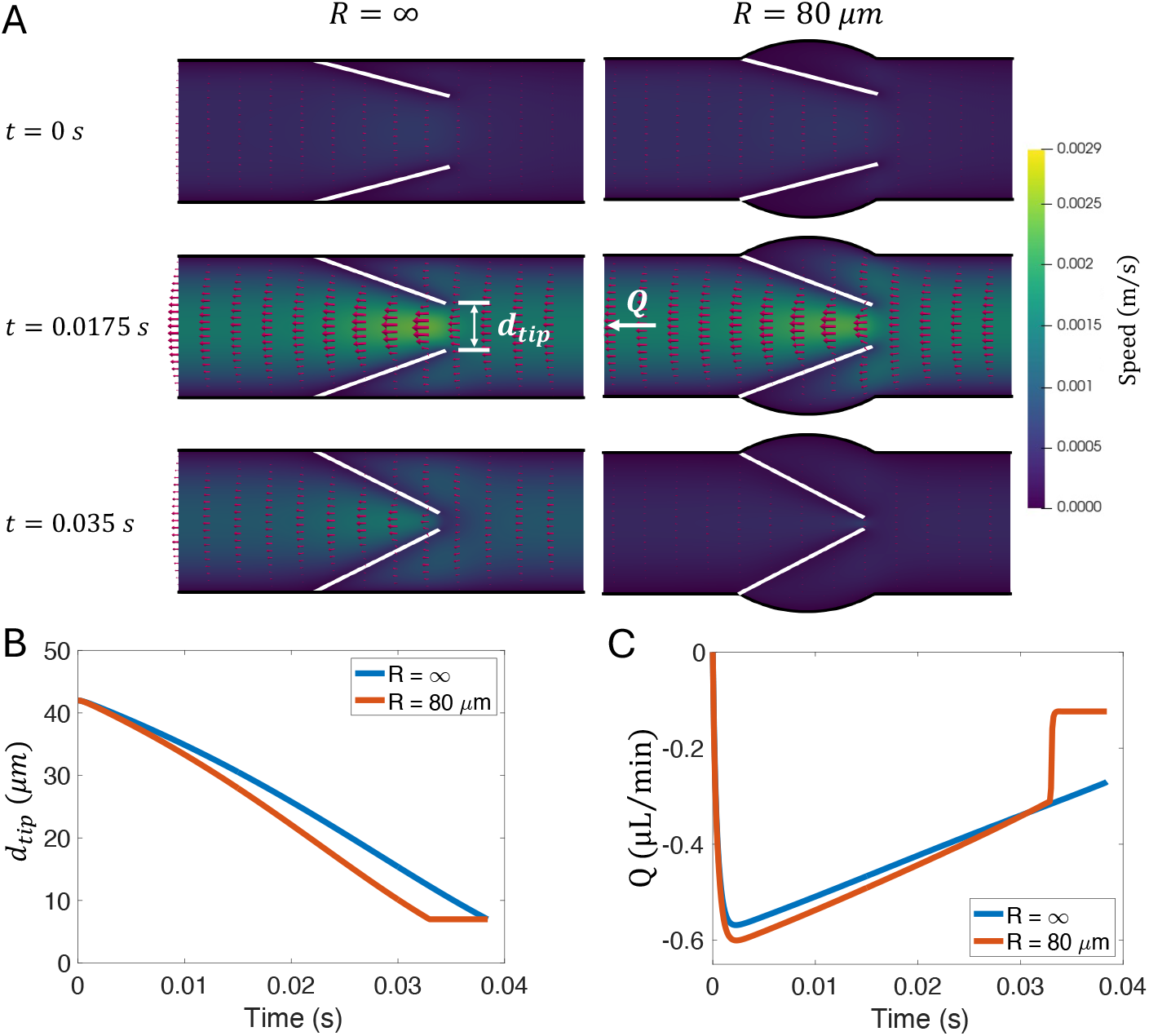
Valve closing dynamics under a constant -1 Pa negative pressure drop, comparing a straight sinus (*R* = ∞) with a circular sinus (*R* = 80 *µ*m). (A) Snapshots of the velocity field, showing backflow (leftward) at *t* = 0, 0.0175, and 0.035 s. Distance between valve tips (*d*_*tip*_) in time series for the straight sinus (blue curve) and circular sinus (orange curve). Time series of instantaneous upstream volumetric flow rate (*Q*_*upstream*_), or backflow for both cases.

It was found that the valve with circular sinus closed noticeably faster than the straight sinus, as seen in Figure 6B. This plot of the valve tip distance (*d*_*tip*_) shows the circular sinus valve (orange curve) fully closing at *t* ≈ 0.032 s, while the valve in the straight sinus (blue curve) was still in motion.

An interesting phenomenon was observed in the upstream volumetric flow rate (backflow), shown in Figure 6C. Because the circular sinus valve closes faster, its motion creates a slightly larger initial backflow than the straight sinus case. However, this backflow is suddenly reduced at *t* ≈ 0.032 s as the valve shut, settling to a small, constant leakage (around -0.08 *µ*L/min). In contrast, the slower-closing straight sinus valve allows a larger backflow to persist for a longer duration. This difference in total backflow was quantified by computing the time-averaged backflow 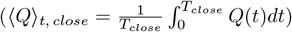 . It was found that 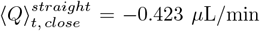 versus 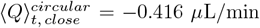. The total backflow in the straight sinus case was 1.65% larger than in the circular sinus case. This demonstrates that the circular sinus geometry not only enhances valve opening but also promotes a faster, more effective closure that is more efficient at blocking backflow.

## 4 Discussion

In this study, we developed a 2D fluid-structure interaction model to investigate cerebrospinal fluid (CSF) drainage through a cervical lymphatic vessel (CLV). We confirmed that the system is governed by a complex interplay of active wall pumping and passive valve dynamics, evidenced by the phase lag observed between the wall motion and the resulting pressure peaks. In addition, we quantified the importance of inter-lymphangion coordination, showing that a prograde propagation of contraction is more effective at driving flow than synchronous or retrograde propagation of contractions. Our results also demonstrate that the circular valve sinus, evident in histological images [16, 17], is more efficient than a straight sinus, enhancing both opening and closing dynamics to improve net flow.

Based on our single-lymphangion model, the estimated Reynolds number 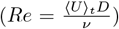 is approximately 0.0067 and the Womersley number 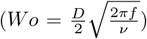 is approximately 0.047. Both of these are close to measurements in physiological conditions reported in Du et al (*Re* = 0.0064 ± 0.0008 and *Wo* = 0.039 ± 0.003 respectively) [8]. These values confirm the flow is in the creeping flow regime, where viscous forces are dominant.

Our simulation results from the single-lymphangion model suggest that a region of negative relative pressure exists inside the CLV (Figure 3C). This coincides exactly with the time when the lymphangion is in diastole (Figure 3B). This negative pressure was also reported by Jamalian et al. [50], who termed it a *suction* pressure and noted that it allows fluid to be drawn in through the initial lymphatics. This suction is caused by the elastic re-expansion of the lymphangion (i.e., passive elastic recoil and geometric recovery). Although we used prescribed wall motions, our model accurately captures this suction pressure phenomenon.

A key question is whether this suction pressure is strong enough to extract CSF from the subarachnoid space. The connection between the subarachnoid space and the CLV is mediated by meningeal lymphatics [13] and nasal lymphatics within the nasopharyngeal lymphatic plexus [16]. These newly discovered vessels are reported to lack smooth muscle cells, meaning they likely do not pump. Therefore, for CSF to be withdrawn purely by the recoil of a CLV, the suction pressure would have to be strong enough to overcome the hydraulic resistance of the complicated and long connection bridging the CLV and the subarachnoid space. Further experimental work is required to elucidate the clear map of the CSF drainage route and its primary driving forces.

We quantified the volumetric flow rate at the downstream outlet for various inter-lymphangion contraction phase delays, ultimately to test which pattern is the most effective. A central finding of our study is the superior efficiency of the prograde propagation of the contraction wave with a *π/*2 phase delay. This suggests that the Δ*Φ* = *π/*2 delay creates an optimal peristaltic-like wave, which effectively propels the fluid forward while reducing the backflow seen in the more abrupt synchronous or *π/*4 contractions. This result highlights a key trade-off: while the synchronous and *π/*4 delay contraction cases generate a higher instantaneous peak flow, they also induce a larger backflow, ultimately reducing their net efficiency. The *π/*2 delay appears to achieve the optimal balance between a strong forward flow and minimized reflux.

Our result that a prograde contraction propagation wave is optimal does not perfectly match the findings from Elich et al. [29], which was based on rat mesenteric lymphatics. They reported that synchronous contraction of lymphangions yielded the highest volumetric flow rate, although we both agree that retrograde propagation results in the least efficient pumping. This discrepancy may arise from key differences in our model formulations. For instance, our valve sinuses are fixed, while their sinus slightly moved with the vessel wall contraction. Most importantly, our model uses a significantly higher aspect ratio (*L*_*lymphangion*_*/D*_*lymphangion*_ = 4.76) compared to the ratio of 3.2 used in Elich et al. [29]. This higher aspect ratio is a deliberate choice based on recent CLV imaging [8, 12, 16, 17], as CLVs are observed to be characteristically slender. This key geometric difference, which better reflects the in vivo morphology, may contribute to the different pumping dynamics observed in our results.

It is important to note that *Q* in the three-lymphangion test was computed just past the third lymphangion (Figure 4A). The highly pulsatile flow seen in Figure 4B is therefore logically dominated by the action of that third, nearest lymphangion. However, a more subtle effect is also visible. In the prograde contraction plots (e.g., the orange curve, Δ*Φ* = *π/*2, from *t* = 0 − 2.5 s), small increases in *Q* appears before the third lymphangion contracts. These are the flow rates generated by the first and second lymphangions, indicating our model captures their upstream contribution. However, these upstream effects are minor. This suggests that in the extremely low Reynolds number regime of the flow in the CLVs, these propagated pressure pulses are rapidly damped by the high hydraulic resistance as well as the compliance of the flexible membranes and valve motions. Consequently, the contraction of a distant lymphangion has a weak influence on the local flow. This implies that flow dynamics observed in vivo, which are often limited to a small field of view [4], are likely dominated by local, not global, pumping events *if* no strong external pressure is applied.

We conducted a separate valve test to quantify the hydrodynamic efficiency of the straight sinus versus the circular sinus. We found that the circular sinus is more effective at both promoting forward flow during opening and blocking backflow during closure. This finding aligns well with the numerical study by Wilson et al. [30]. Although their model differed in geometry, was 3D, and was not a fully-coupled FSI simulation, they similarly reported that the circular sinus reduces friction and thus lowers the overall hydraulic resistance. We extended this analysis by also testing the closing dynamics. Our results revealed that the faster motion of the circular-sinus valve itself creates a slightly larger instantaneous backflow over much of the closing cycle (Figure 6C). However, because the valve shuts much faster, the total backflow is smaller compared to the straight sinus, which allows a slower, persistent retrograde leak for a longer duration.

Quantitatively, the time-averaged forward flow during opening (Figure 5) was 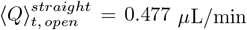 and 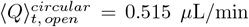 . During closure (Figure 6), the time-averaged backflow was 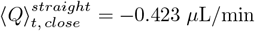 and 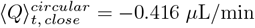 . While the resulting differences of 7.97% (opening) and 1.65% (closing) may seem small and negligible, this test was for an isolated component of a single valve. In a physiological system, where hundreds or thousands of valves are connected in series and parallel through the CLV [4], meningeal [13, 51, 52], and nasopharyngeal [16] networks, these seemingly minor local efficiencies would likely compound into a significant, system-level improvement in overall transport performance.

There are several limitations to be noted in this study. First, the secondary valves were modeled as rigid bodies rotating about a hinge. In reality, these leaflets posses a distinct curvature and are flexible [8]. A future direction may include implementing fully deformable, elastic leaflets, potentially by coupling the computational fluid dynamics solver with a finite element analysis, instead of IB formulations, to model the true leaflet mechanics. This would also address inherent limitations of the IB, which approximates the no-slip condition, resulting in small amounts of fluid leakage across the wall [53, 54]. However, FSI simulations coupled with finite element analysis are computationally expensive, and the specific mechanical properties of CLV components remain to be fully characterized. A second limitation is the non-zero resting gap (*d*_*tip*_) in our model, which means the valve is never fully closed. While the valves are biased open under typical resting conditions [37], they likely fully close under a strong adverse pressure difference across the valve. This persistent gap, combined with the zero-pressure boundary conditions, is the primary reason for the low-pressure range observed in Figure 3C compared to 0D models [19, 21, 55]. It is important to note, however, that this pressure range is consistent with other 2D simulations [27]. Unfortunately, in vivo pressure measurements from murine CLVs would be experimentally difficult to obtain for validation.

A further limitation is that this study employed a 2D simulation framework. To estimate the volumetric flow rate, *Q*, we first computed the flux by integrating the streamwise velocity component across the channel width and then multiplied by the lymphangion diameter, assuming the velocity is uniform in the out-of-plane direction, similar to prior 2D studies [27, 29]. This approximation does not fully capture the fluid dynamics of real lymphatic vessels, whose cross-sections are circular. In addition, lymphatic valves are bicuspid, 3D structures [31, 37]. Although the low Reynolds number confirms a creeping flow regime [8], the complex 3D shape of the valve and its oval-shape orifice would likely result in different streamlines and velocity fields than our 2D simulation. Expanding this current model to 3D is a logical next step.

Across all our analyses, no significant secondary flow or vortices were observed, which contrasts with other lymphatic simulation studies [29, 31]. This is likely a consequence of the creeping regime in our model. However, it remains unclear if recirculation zones are truly present in CLVs in vivo or if they are a feature of the relatively higher Reynolds number flows in the rat mesenteric lymphatics. Recent in vivo measurements that utilize microsphere injections do not exhibit recirculation behind valves [4, 8], but further high-resolution experimental analysis may be needed.

Finally, we used zero-pressure boundary conditions to isolate the flow driven purely by the intrinsic pumping and valve dynamics. In reality, the upstream pressure may be linked to the intracranial pressure [52] and a complex upstream network, possibly even including the recently identified arachnoid cuff exits [56]. As new experimental data emerges revealing anatomical details of this network, future simulations could implement a multi-scale model. Coupling the 3D fluid-structure model with a lumped-parameter model (similar to Windkessel boundary models in the cardiovascular system) representing the full upstream network may help to better understand the CSF drainage dynamics and potentially lead to therapeutic strategies for restoring/maintaining CSF drainage.

## 5 Conclusions

In this study, we developed a fully-couple 2D fluid-structure interaction (FSI) model of murine cervical lymphatic vessels (CLV), parameterized by recent in vivo measurements, to investigate the cerebrospinal fluid (CSF) drainage dynamics along the cervical lymphatic pathway. Simulations of a single lymphangion successfully captured the fundamental pumping mechanism, demonstrating how wall contractions and expansions generate the pressure gradients that drive the flow. Expanding the model to a three-lymphangion chain revealed that the net transport critically depends on the phase delay between contractions, with an prograde propagation wave being vastly more effective than retrograde patterns. Finally, we elucidated the influence of the valve sinus geometry, showing that circular sinuses enhance both effective valve opening and closure, thereby promoting net flow.

To our knowledge, this is the first detailed FSI simulation to explicitly model the murine CLV system, establishing a new mechanical framework for understanding this CSF drainage pathway. By bridging in vivo anatomical and physiological observations with computational modeling, this framework provides a mechanistic basis for understanding how CLV contractility and valvular dynamics regulate CSF outflow from the brain. These findings may advance the broader understanding of impaired CSF outflow in aging, after traumatic brain injury, or in many other neurological disorders, offering new insights for designing future experimental studies and developing therapeutic strategies.

## 6 Availability of data and materials

The simulation used in this study is available from the corresponding authors upon reasonable request.

## 8 Author information

### Contributions

Conceptualization: DK, JT; Methodology: DK, JT; Formal analysis and investigation: DK; Writing - original draft preparation: DK; Writing - review and editing: JT; Funding acquisition: JT; Resources: JT; Supervision: JT

## 9 Ethics declarations

### Ethical Approval

Not applicable.

### Consent for publication

Not applicable.

## Appendix A Validation of Lattice Boltzmann method coupled with Immersed Boundary solver

**Fig. A1.**
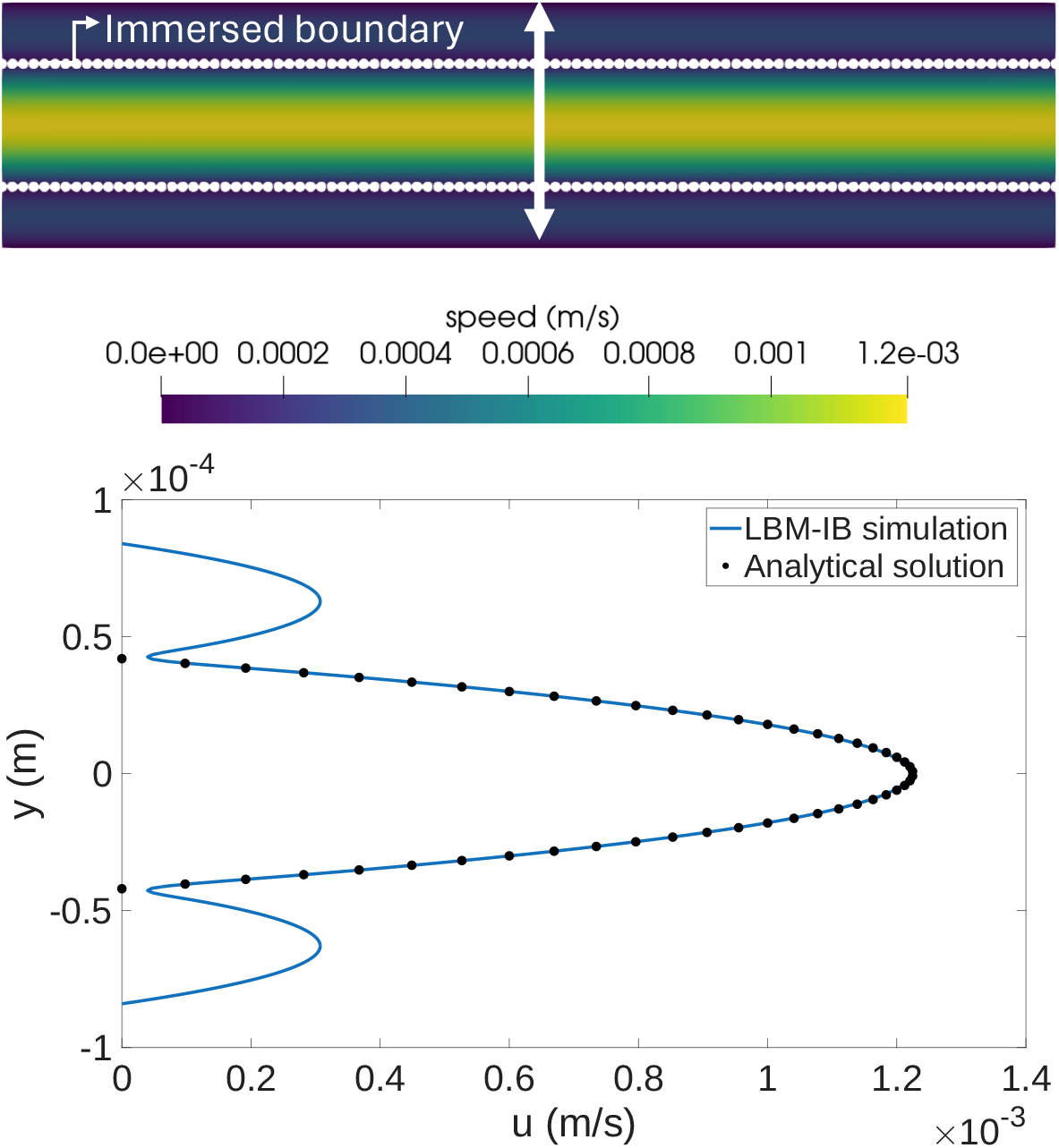
A1. Verification of the Lattice Boltzmann method coupled with Immersed boundary (LBM-IB) solver for 2D Poiseuille flow. The velocity profile from the LBM-IB simulation (solid line) is compared against the analytical solution (dots).

To verify the coupled Lattice Boltzmann (LBM) and Immersed Boundary (IB) solver, it was validated against the analytical solution for Poiseuille flow, which describes pressure-driven flow between two stationary parallel plates. Given a pressure drop Δ*P* over a length *L*, the parabolic velocity profile *u*(*y*)

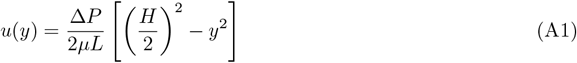

where *H* is the channel height and *µ* is the viscosity of the fluid.

This analytical solution was compared against the LBM-IB simulation, where we placed static Lagrangian points within the domain to mimic a no-slip wall (labeled as ‘Immersed boundary’ in Figure A1) using:

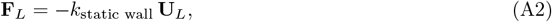

where the description of the constants and variables are explicitly explained in the main text.

The pressure-driven flow was implemented using the Zou-He boundary condition. At the inlet (left wall), a pressure *P*_*in*_ = 1 Pa is set by fixing the density *ρ*_*in*_. The unknown velocity *u*_*x*_ is first computed:

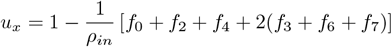

Then, the unknown incoming populations (*f*_1_, *f*_5_, *f*_8_) are calculated:

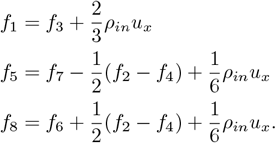

A similar formulation was applied at the outlet (right wall) with *P*_*out*_ = 0 Pa. The top and bottom walls of the fluid domain were treated with the standard on-grid bounce-back scheme, where incoming populations are reflected to the opposite direction. Specifically, *f*_2_ = *f*_4_, *f*_5_ = *f*_7_, and *f*_6_ = *f*_8_ were used for the bottom wall, and *f*_4_ = *f*_2_, *f*_7_ = *f*_5_, and *f*_8_ = *f*_6_ were used for the top wall. A detailed definition, derivation, and implementation can be found in the LBM textbook that served as the basis for this work [39]. As seen in Figure A1, the LBM-IB solver matches well with the analytical solution. The difference in the maximum x-velocity (*u*_*max*_) between the analytical solution and the LBM-IB simulation was 0.3%.

## Appendix B Grid Convergence Test

A grid convergence study was conducted to ensure that the numerical results are independent of the spatial discretizations of the fluid domain. The simulation of a single lymphangion was used as the test case. Several simulations were performed with increasingly finer grid resolutions, while all Immersed boundary parameters were held constant.

Figure B2A shows ⟨*Q*_*out*_⟩_*t*_ as a function of the total number of grid cells. As the grid is refined, the calculated flow rate increases and begins to plateau, asymptotically approaching a grid-independent value. Figure B2B provides a more quantitative look at the convergence. It shows the relative error of ⟨*Q*_*out*_⟩_*t*_ for each grid, using the solution from the finest grid (94,560 nodes) as the benchmark. As the grid size decreases (i.e., resolution increases), the relative error decreases.

**Fig. B2.**
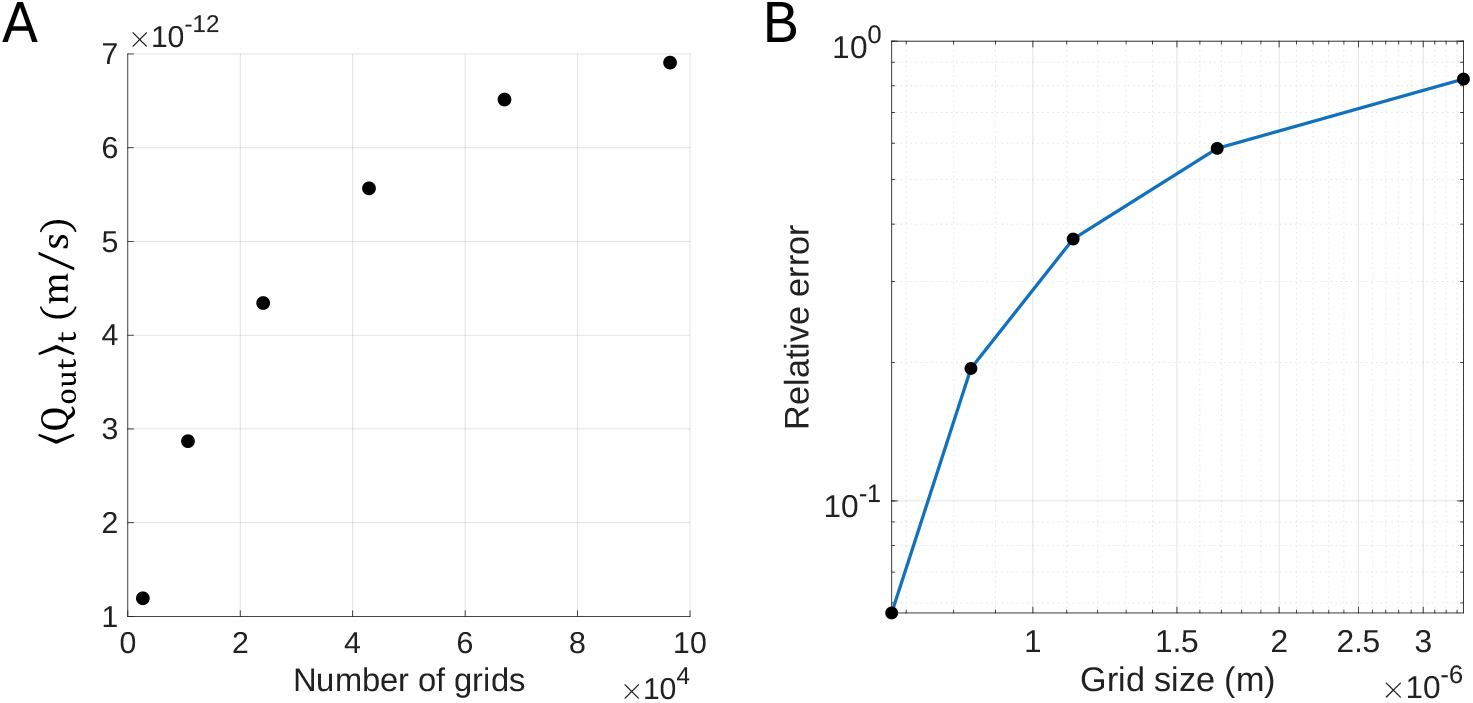
. Grid convergence study for the single lymphangion simulation. (A) The time-averaged outlet flow rate (⟨*Q*_*out*_ ⟩_*t*_) is plotted against the total number of fluid grid nodes. (B) The relative error for each grid resolution, calculated with respect to the finest grid (94,560 cells), is plotted against the grid size.

For the two finest grid tested (67,000 and 94,560 cells respectively), the relative error in ⟨*Q*_*out*_⟩_*t*_ between them was calculated to be approximately 5.7 %. Given that the computational cost increase exponentially with finer discretizations, a balance between solution accuracy and computational efficiency must be chosen. Therefore, the 67,000-cell grid was selected for all subsequent simulations, as it provides a reasonably accurate solution at a feasible computational cost.

